# Synthesis of natural history collections data reveals patterns of US freshwater mussel diversity and decline

**DOI:** 10.1101/2022.09.22.509037

**Authors:** John Pfeiffer, Traci P. Dubose, Sean M. Keogh

## Abstract

Natural history collections are uniquely positioned to chronicle biodiversity changes across time and space and are a fundamental data source in taxon-based research and conservation. With over 90 species listed under the Endangered Species Act, freshwater mussels are one of the most imperiled animal assemblages in the United States and are the focus of considerable conservation efforts (e.g., species status assessments, listing decisions, and recovery plans). Unfortunately, natural history collections data is often underleveraged in such efforts, in part, because much of the data are decentralized and nonstandard, and thus, difficult to access and analyze. Our objective herein is to synthesize, standardize, and enrich digitized US freshwater mussel collections data to better suit the needs of conservation stakeholders. We aggregated specimen records from 45 US natural history collections and enriched these records by programmatically standardizing taxonomic information, flagging potentially problematic records, and joining records with freshwater-specific spatial frameworks (e.g., hydrological units and stream segments) and their associated metadata (e.g., area, stream order, discharge, velocity). The assembled dataset includes 410,665 records, 302 species, and 1,494 hydrological units (8 digit-level). Using these enriched records, we estimated ecological attributes for over 280 freshwater mussel species including aspects of range size (i.e., area of occupancy and change in area of occupancy) and habitat preferences (i.e., stream order size, discharge, slope, and velocity). Listed species had significantly fewer occurrences (p<0.001) and smaller area of occupancy (p<0.001) in comparison to non-listed listed species. Listed species also tended to have a higher stream order preference (p<0.001) and discharge preference than non-listed species (p<0.001). These important ecological attributes have not been incorporated into freshwater mussel conservation efforts in a quantitative way and our novel estimates can be used to make more data-driven ecological and conservation inferences.

## Introduction

Specimen-based occurrence records are a basic yet powerful tool to understand the natural history of species including how spatial and ecological patterns have changed over time — a critical perspective given the escalating biodiversity crisis and the increasing importance of taxon-based conservation. Natural history collections provide the physical and digital infrastructures that link voucher specimens to their associated spatial and temporal data, which allows occurrence data to dynamically change with ever-growing understanding of biodiversity. Digitization and aggregation of natural history collections data has grown dramatically in recent decades and has helped to usher biodiversity science into the big data era (Nelson & Ellis 2019; Hedrick et al. 2020). However, much digitization and aggregation remain to be done, and for rapidly declining assemblages like the native freshwater mussels (Unionidae + Margaritiferidae) of the United States, there is an urgent need for a more comprehensive synthesis of occurrence records to better document the distribution, ecology, and trends of imperiled species.

Ninety of the 303 freshwater mussel species native to the United States are listed as Threatened or Endangered under the Endangered Species Act (ESA) (50 CFR § 17.11; Williams et al. 2017). That is, 30% of US freshwater mussel species are in danger of extinction according to federal law, with an additional 34 freshwater mussel species being petitioned or proposed for listing under the ESA (USFWS Species Data Explorer - https://ecos.fws.gov/ecp/report). The dramatic decline of US freshwater mussel diversity, distribution, and abundance is well-documented and has relied, in part, on natural history collections data (e.g., Haag 2019; Smith et al. 2019; Pursifull et al. 2021). The abundance of US freshwater mussel specimens and their associated occurrence data (e.g., identification, locality, date, etc.) deposited in natural history collections provide the permanent vouchers necessary to verifiably measure how mussel biodiversity has changed across time and space. Despite the value of natural history collections in freshwater mussel research and conservation, these institutions and the data they curate remain underleveraged.

One of the factors limiting greater use of natural history collections data is that much of the digitized data is decentralized and nonstandard. While many US freshwater mussel collections have been ingested by biodiversity aggregators like Global Biodiversity Information Facility (www.GBIF.org) and InvertEBase (www.invertebase.org),, many important collections are not yet included in those infrastructures. Consequently, there is no comprehensive, centralized, and easily accessible inventory of digitized US freshwater mussel occurrence records. Given the alarmingly high levels of imperilment, there is an urgent need for a more synthetic inventory of existing freshwater mussel collections data customized to better meet the needs of conservation stakeholders. Our objectives are 5-fold:

1. Create an inventory of digitized, US freshwater mussel collections data by synthesizing, standardizing, and programmatically validating specimen-based occurrence records from 45 US natural history museums;
2. Enrich occurrence records by linking them with freshwater spatial frameworks and their associated hydrological variables (e.g., hydrologic units, stream order, discharge, velocity, slope);
3. Examine how occurrence data varies across institution, mussel biodiversity, time, and space;

4.Use this resource to estimate species specific ecological attributes (e.g., range size and habitat preference);

5.Translate findings into data-driven recommendations about how to better use and improve freshwater mussel natural history collections data.

## Methods

### Standardization of Digitized US Freshwater Mussel Occurrences

We obtained digitized specimen-based occurrences of freshwater mussels (Unionidae + Margaritiferidae) from the United States from 45 US natural history collections by requesting records directly from collection managers and curators. The following variables were included in our inventory: institution, catalog number, taxonomic identification, country, state, locality, latitude, longitude, day, month, year, collector, remarks, and preparation. Occurrence records were then aggregated and values for country, state, month, day, and year were standardized using OpenRefine (https://openrefine.org/). Day, month, or year values given as a range (e.g., pre-1900, Spring) were treated as missing data. All subsequent data manipulation and analysis was done in R (R Core Team 2022). All files and scripts necessary to reproduce the dataset and analyses can be found in the Supplementary Information (Appendices S1-3).

Georeferences were standardized by converting all coordinates to decimal degrees. We assumed that all US records should be in the northwestern hemisphere and adjusted the signs of coordinates accordingly (latitude > 0, longitude < 0). We assigned all georeferenced records to USGS Hydrological Unit Codes at the 2-digit through 10-digit levels (HUC2-HUC10) within the Watershed Boundary Dataset spatial framework using the sf R package (Pebesma 2018; USGS 2020). Georeferenced records plotted outside the Watershed Boundary Dataset were flagged and excluded from downstream analyses. We verified that georeferenced locality was within 10 kilometers of the Albers projection of the recorded state value; if not, the record was flagged and excluded from downstream analyses. To identify the potential flowline segment (COMID) represented in each record, we first removed flowlines with stream order equal to 1 from the National Hydrography Dataset Plus Version 2 (NHDPlusV2; McKay et al. 2012) to reduce the likelihood of incorrectly pairing an occurrence with a flowline given potentially imprecise georeferences and the abundance of 1^st^ order streams. We then spatially joined (snapped) each occurrence to the closest flowline segment and recorded the COMID.

Taxonomic identification was standardized using the GBIF Taxonomic Backbone (GBIF Secretariat 2021) via the R function *rgbif*::*name_backbone* (Chamberlain et al. 2017). Some identifications were being matched to a higher taxonomic rank (e.g., genus) due to misspellings or unusual taxonomic combinations. We created a refined_name variable by duplicating the verbatim_name variable and making minor changes to those names so they matched more precisely when re-running the *name_backbone* function. The resulting dataset was filtered to include records from native US families, genera, and species as recognized by the Freshwater Mollusk Conservation Society’s Scientific and Common Names Committee (FMCS 2021). The GBIF taxonomic backbone is mostly congruent with the FMCS Scientific and Common Names Committee, but some minor changes were made (e.g., *Toxolasma pullum* to *Toxolasma pullus, Obovaria jacksoniana* to *Obovaria arkansasensis*). All taxonomic modifications are documented in Appendix S1.

Over 70 North American freshwater taxa have been either described or elevated from synonymy since 2007 (Graf & Cummings 2021) and, as a consequence, the identification of many occurrence records are in need of updating. We did a literature review of taxonomic revisions over the last twenty years and identified 17 species that required updating. In situations where a widespread species was revised to multiple, allopatric species, we updated identifications based on location to reflect these recent taxonomic revisions using the R package *CoordinateCleaner* (Zizka et al. 2019). For all 303 species, we did a literature review, created a range map shapefile based on aggregation of HUC10’s, and flagged records that occurred outside the shapefile boundary. Our script greatly benefited from existing code at https://github.com/dmacguigan/USHiddenRivers.

Typically, freshwater mussel collections have digital records for a single taxon with identical spatial and temporal data (i.e., a “lot” in most mollusc collections). Occasionally, institutions will have digital records for every individual which duplicates occurrences and makes comparisons difficult. To standardize data for downstream analyses we used occurrences which is defined here as a single record at an institution with identical taxonomic, spatial, and temporal data. We programmatically identified occurrences using the *campfin* R package (https://github.com/irworkshop/campfin) and flagged duplicate records.

### Summarizing Digitized Occurrences

For each natural history collection in the assembled inventory, we calculated the total number of records, occurrences, genera, species, and occupied HUC8s, as well as the percentage of occurrences identified to species, percent with year data, percent georeferenced, and percent stored as dry specimens. We also determined the number flagged occurrences (i.e., duplicate records, state/georeference mismatch, outside Watershed Boundary Dataset, outside species’ expected distribution). These institutional summary statistics and the synthesized collection data were visualized to reveal temporal, taxonomic, and spatial patterns in digitized US freshwater mussel collections data.

### Estimating Ecological Attributes

Non-flagged, georeferenced occurrences identified to the species-level were used to estimate species’ ecological attributes (e.g., area of occupancy, mode stream order, median discharge, etc.). We estimated ecological attributes for species with more than 10 georeferenced occurrences. We estimated a species’ area of occupancy (AOO) by identifying HUC8s that contained at least one georeferenced occurrence and summing the area of the occupied HUCs. We report the minimum and maximum latitude and longitude for each species and used a convex polygon of all georeferenced points to identify the middle latitude and longitude. For each occurrence successfully linked to a flowline segment (COMID), we extracted several hydrological variables from NHDPlusV2: Strahler stream order, slope of flowline (difference in smoothed elevations in meters/stream segment length in meters), mean annual discharge (cubic feet/second; cfs), and mean annual velocity (feet/second). Mean annual discharge and median mean annual velocity represent the best available estimate of stream flow based on the Enhanced Runoff Method; other estimates of annual discharge and velocity, along with monthly estimates, are available in the NHDPlusV2 and can be queried with a COMID (Moore et al. 2019). Nonparametric measurements of central tendency were used to estimate each species’ mode stream order, median slope, median discharge, and median velocity preference.

We also measured how each species’ AOO has changed across time according to specimen-based occurrences. To correct for taxonomically and temporally uneven collection effort, we split each species collection history (i.e., all non-flagged occurrences with year data) by its median collection year which resulted in two time frames with roughly equal number of occurrences. We then measured what proportion of the total AOO was accounted for in the first and second halves of the species collection history, and how the proportion has changed between the time frames. These calculations are designed to determine how the collection of a species has changed over time while accounting for temporal biases in collecting effort and is not necessarily a reflection of changes in the species distribution (Campbell Grant 2015).

### Estimating HUC Attributes

Using non-flagged species-level occurrences, we also estimated the number of occurrences, species richness, and temporal changes in species richness for each HUC8 in the United States. We estimated species richness changes using the same approach as estimating changes in a species AOO, described above. These calculations are designed to determine how the collecting in a HUC has changed over time while accounting for temporal biases in collecting effort and may not necessarily reflect changes in a HUC’s species richness.

A species was considered “listed” if its ESA Listing Status was “Endangered”, “Threatened”, or “Extinction” according to the USFWS Species Data Explorer (https://ecos.fws.gov/ecp/report) accessed on September 13, 2022. Species that are widely considerd to be extinct (Haag 2012) as were also categorized as “listed”. Non-listed species included all other ESA Listing Status categories (i.e., “Proposed Endangerd”, “Proposed Threatened”, “Resolved Taxon”, “Species of Concern”, “Status Undefined”, and “Under Review”) and all unassessed species. We used a Shapiro-Wilk test was used to test for normality and a Mann-Whitney U test to determine significant differences in 1) the number of occurrences between listed and non-listed species, 2) AOO between listed and non-listed species, 3) percent change in AOO between listed and non-listed species, 4) mode stream order preferences between listed and non-listed species, and 5) discharge preferences between listed and non-listed species.

## Results

### Institutional, Temporal, and Taxonomic Results

We assembled digitized, US freshwater mussel collections data from 45 natural history collections which together included 410,665 records and 360,078 unique occurrences (Appendix S2). The number of occurrences per collection varied from 71,837 (OSUM) to 146 (KU), with a median value of 1,809 occurrences. The total number of species per institution varied from 286 (OSUM) to 29 (KU), with a median species diversity of 141. The maximum number of occupied HUC8s from an institution was 1,021 (USNM), the minimum was 0 (12 institutions), and the median was 51. Collections with more occurrences also tended to have greater taxonomic and geographic coverage (Figure 1A). Institutions varied considerably in their data completeness, especially with respect to percentage of occurrences with year data and percentage of georeferenced occurrences (Figure 1B). The full summary statistics for each institution are described in Appendix S2 and includes additional values such as percentage of occurrences as dry specimens, percentage of occurrences identified to the species-level, and percentage of flagged occurrences.

**Figure 1:**
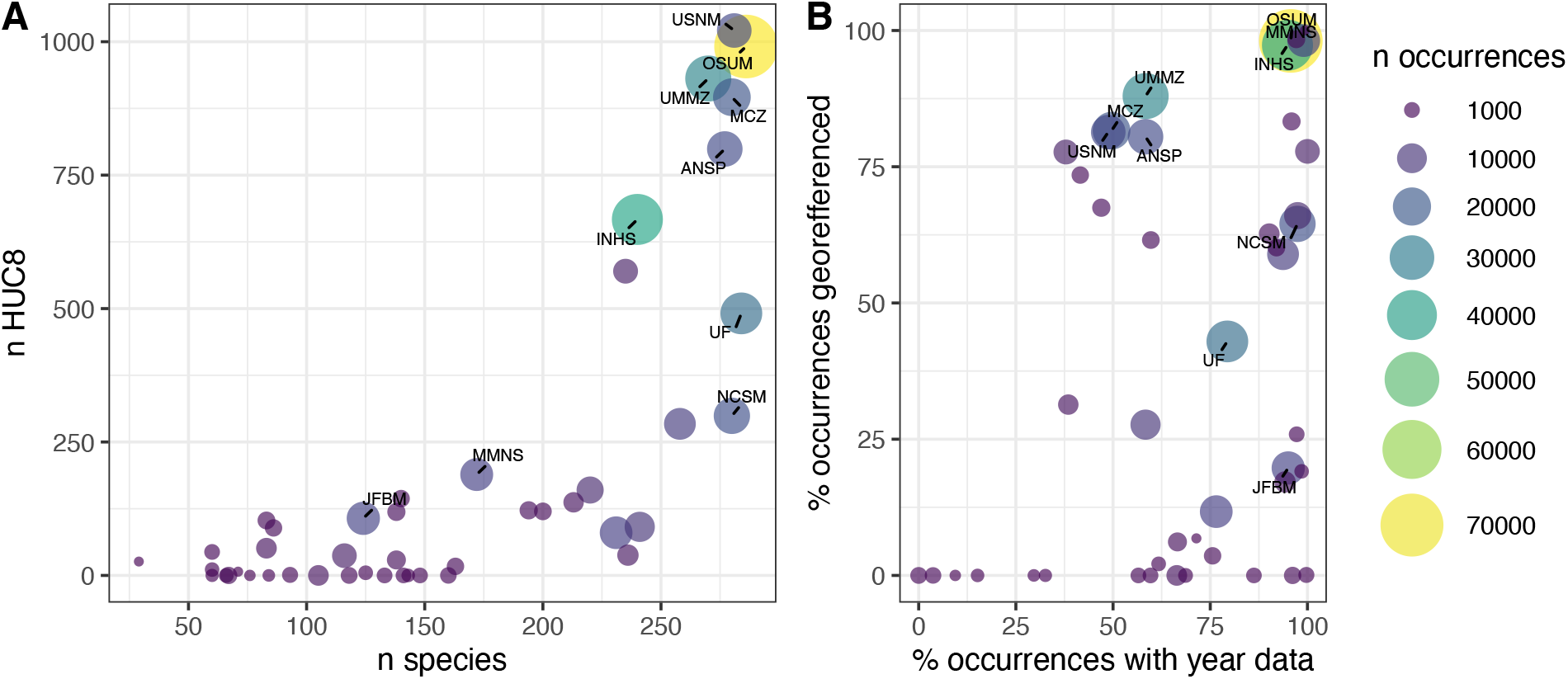
Bivariate plots of the taxonomic and geographic coverage (A) and data completeness (B) of 45 freshwater mussel collections. The ten collections with the greatest number of occurrences are labeled.

Collecting effort is unevenly distributed across time as demonstrated by the 286,280 occurrences with year data (Figure 2). The number of freshwater mussel occurrences per year between 1900-2020 varied from a low of 156 occurrences (1902) to a high of 6,957 occurrences (1987). The mean number of occurrences per year since 1900 is 2,334. The number of occurrences collected per year has been on a downward trend since 2000, and all years since 2011 have been below this 120-year average. This decline is driven primarily by reduced collection of non-listed species. The number of freshwater mussel species collected per year has varied from a low of 54 species (1942) to a high of 234 species (1978). The mean number of species collected per year is 151.5. The number of occupied HUC8’s per year has varied from a low of 32 (1902) to a high of 308 (1976). The mean number of occupied HUC8’s per year is 146.1. The number of species and occupied HUC8s have also been declining since 2000 (Figure 2B-C), but less dramatically than the number of occurrences.

**Figure 2:**
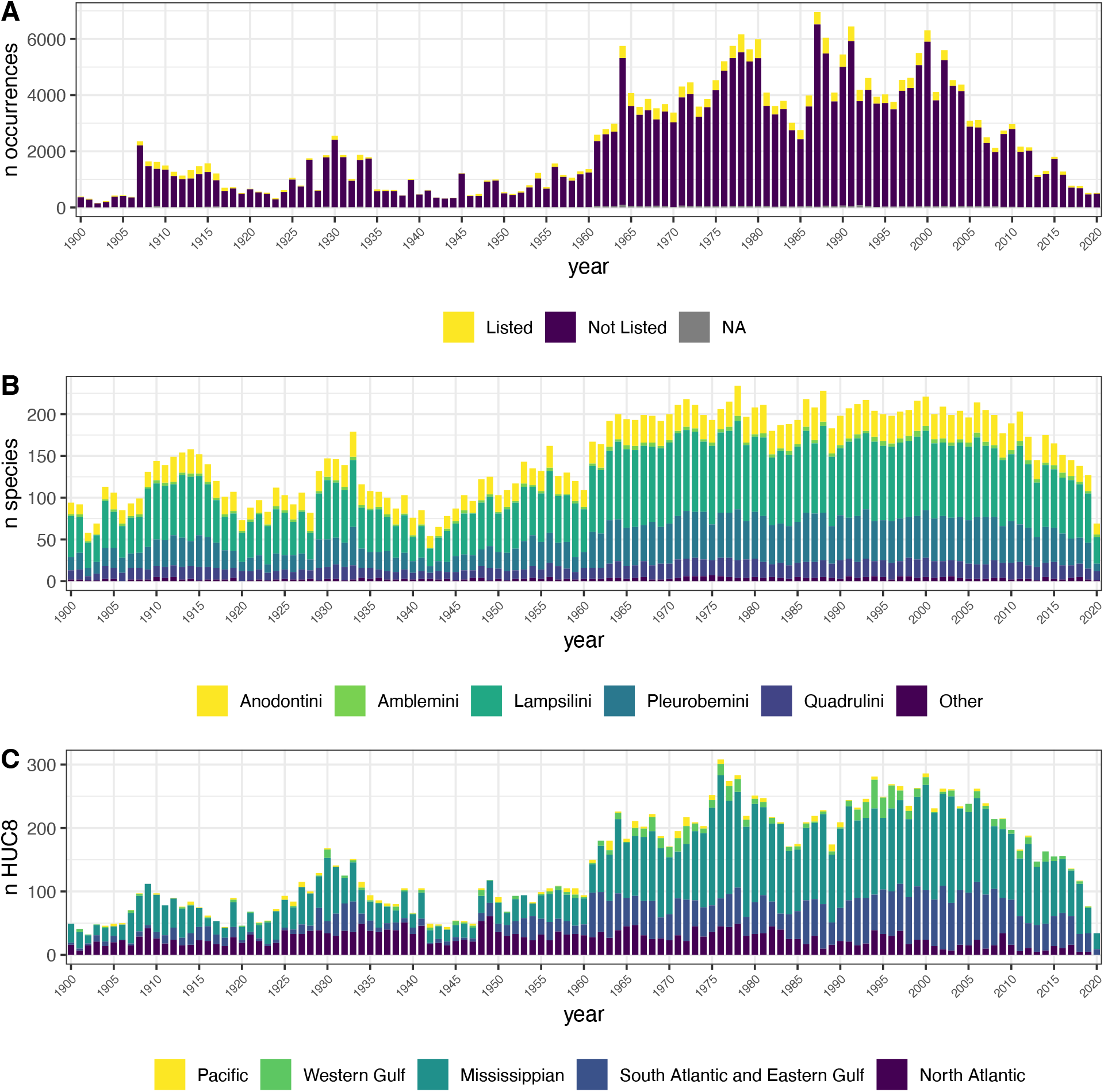
Stacked bar chart of number of occurrences per year (A), number of species-occurrences per year (B), and number of HUC8-occurrences per year (C).

Using the GBIF Taxonomic Backbone, 4,961 unique taxonomic identifications were standardized to 2,014 scientific names. 95.7% of records were assigned via an exact match in the GBIF Backbone, while 2.5% and 1.8% were assigned to a higher taxonomic rank or were fuzzy matched, respectively. Confidence in these assignments had an average of 98.1% (range 80-100).75.4% of the records belonged to accepted names, while 24.5% and <0.001% belonged to synonyms or doubtful names, respectively. Taxonomic precision was high; of the 360,0078 total occurrences, 98.8% are identified to at least the species-level. Occurrences identified to the genus-level were most common in the genera *Elliptio* (1079), *Lampsilis* (463), *Villosa* (285), *Toxolasma* (284), and *Pleurobema* (277). Identification accuracy was also high; of the 253,428 georeferenced species-level occurrences, 95.8% were recovered from their expected range. The number of occurrences per species varied from 1 (*Alasmidonta mccordi*) to 13,752 (*Pyganodon grandis*), with a median value of 341 occurrences per species (Figure 3).

**Figure 3:**
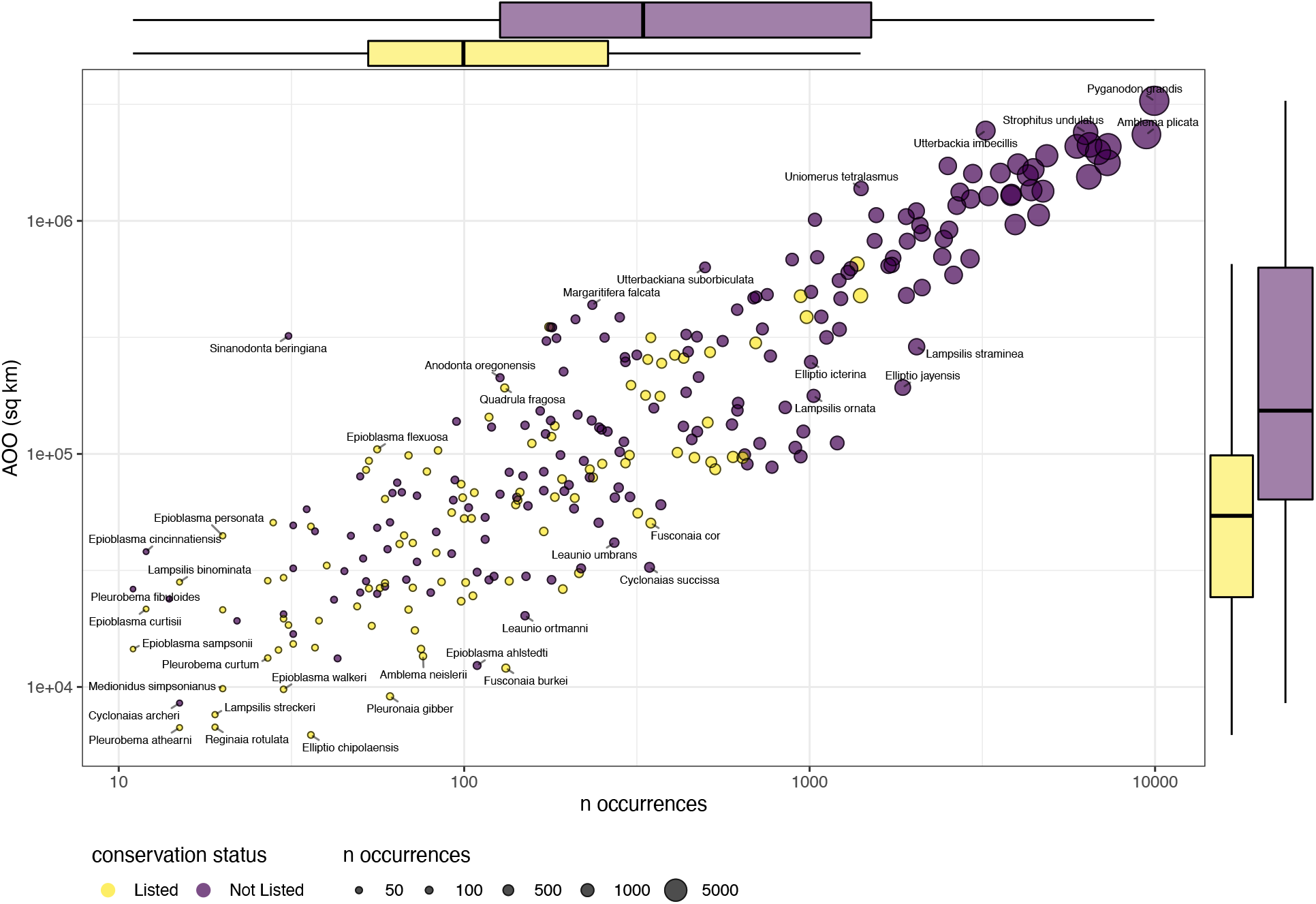
Bivariate plot of number of occurrences (log_10_ scale) and area of occupancy (AOO) with marginal box plots describing variation within listed and non-listed species. Both axes are on the log_10_ scale.

### Ecological and HUC attributes

Of the 302 mussel species in our dataset, 282 species had more than 10 non-flagged occurrences and were used to estimate species’ ecological attributes. *Sinanodonta beringiana* met this threshold but several attributes could not be estimated because NHDPlusV2 flowlines are unavailable in the only US state in which the species occurs (Alaska). Estimates of species’ AOO at the HUC8 level varied from 6,225 km^2^ (*Elliptio chipolaensis)* to 3,276,080 km^2^ (*Pyganodon grandis*) and listed species had significantly smaller median AOO than non-listed species (W = 4,543.5, p-value < 0.001)(Figure 3). Listed species also had significantly fewer median occurrences than non-listed species (W = 4,797, p-value < 0.001). Estimates of AOO change over a species collection history varied from an increase of 70.6% (*Ellipto fraterna*) to a decrease of 75.3% (*Epioblasma ahlstedti*) and the variation between species decreased dramatically with the number of occurrences (Figure 4). Listed species had marginally greater median decreases in AOO in comparison to non-listed species but that result was not statistically different (W = 8,377, p-value = 0.27) (Figure 4). However, when taxa with less than 100 occurrences were excluded from this analysis the result was statistically significant (W = 2,775, p-value = 0.01). Estimates of a species mode stream order varied from 2 (e.g., *Margaritifera margaritifera* and *Lasmigona holstonia*) to 8 (e.g., *Potamilus capax* and *Reginaia ebenus*) and listed species had significantly higher mode stream order preferences than non-listed species (W = 11504, p-value < 0.001)(Figure 5). Estimates of a species’ discharge preference varied from 6.96 cfs (*Margaritifera hembeli*) to 39,203 cfs (*Lampsilis higginsii*) and listed species had significantly higher median discharge preferences than non-listed species (W = 11968, p-value < 0.001)(Figure 5). Summary statistics of all estimated ecological attributes are available in Appendix S2 and should be interpreted using nonparametric statistics and at an ordinal scale. Species summaries can also be accessed via the web app ‘MusselMapR’ (https://musselmapr.shinyapps.io/hic_sunt_naiades/) developed with the R package *shiny* (Chang et al. 2022).

**Figure 4:**
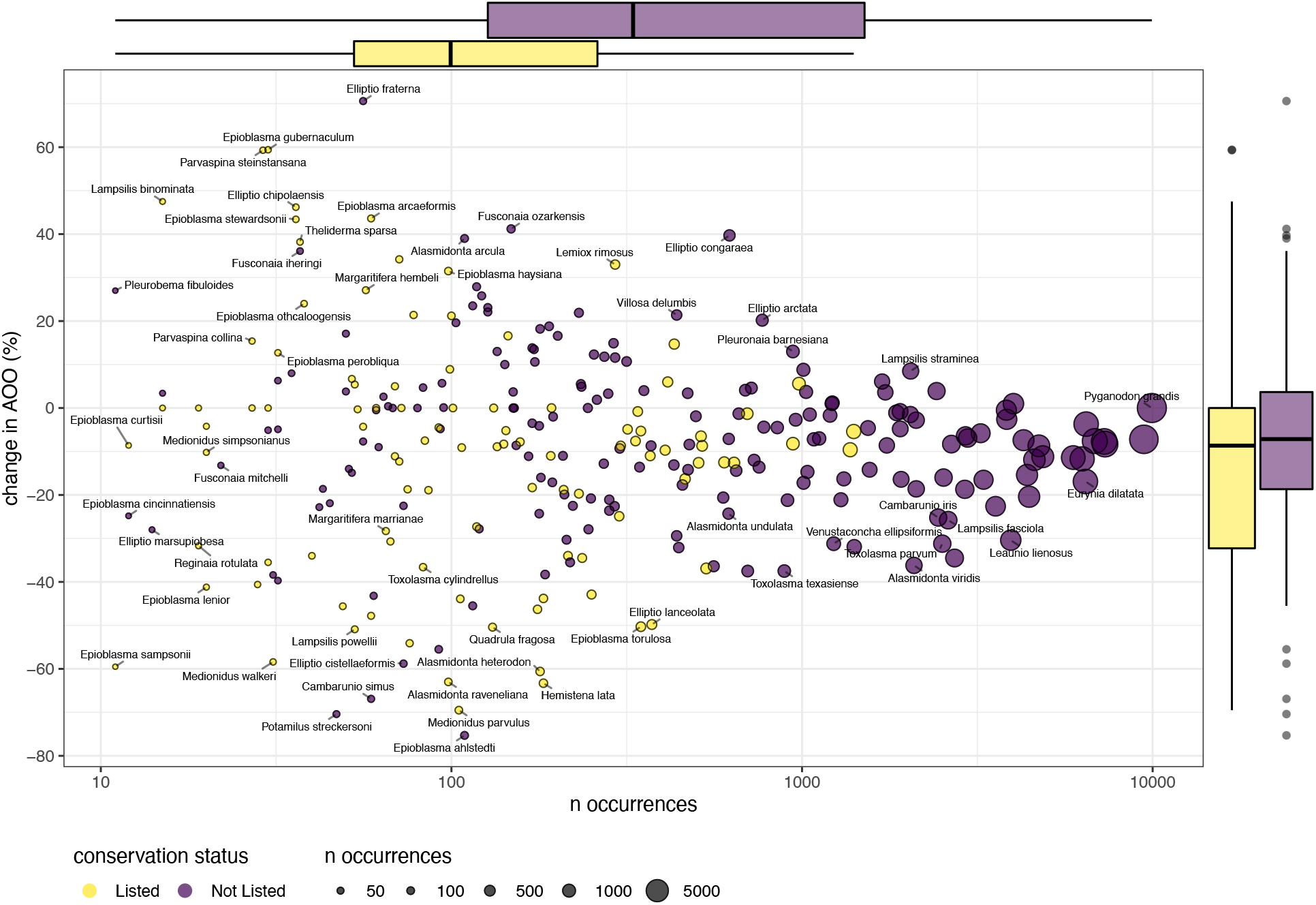
Bivariate plot of number of occurrences (log_10_ scale) and change in area of occupancy (AOO) with marginal box plots describing variation within listed and non-listed species.

**Figure 5:**
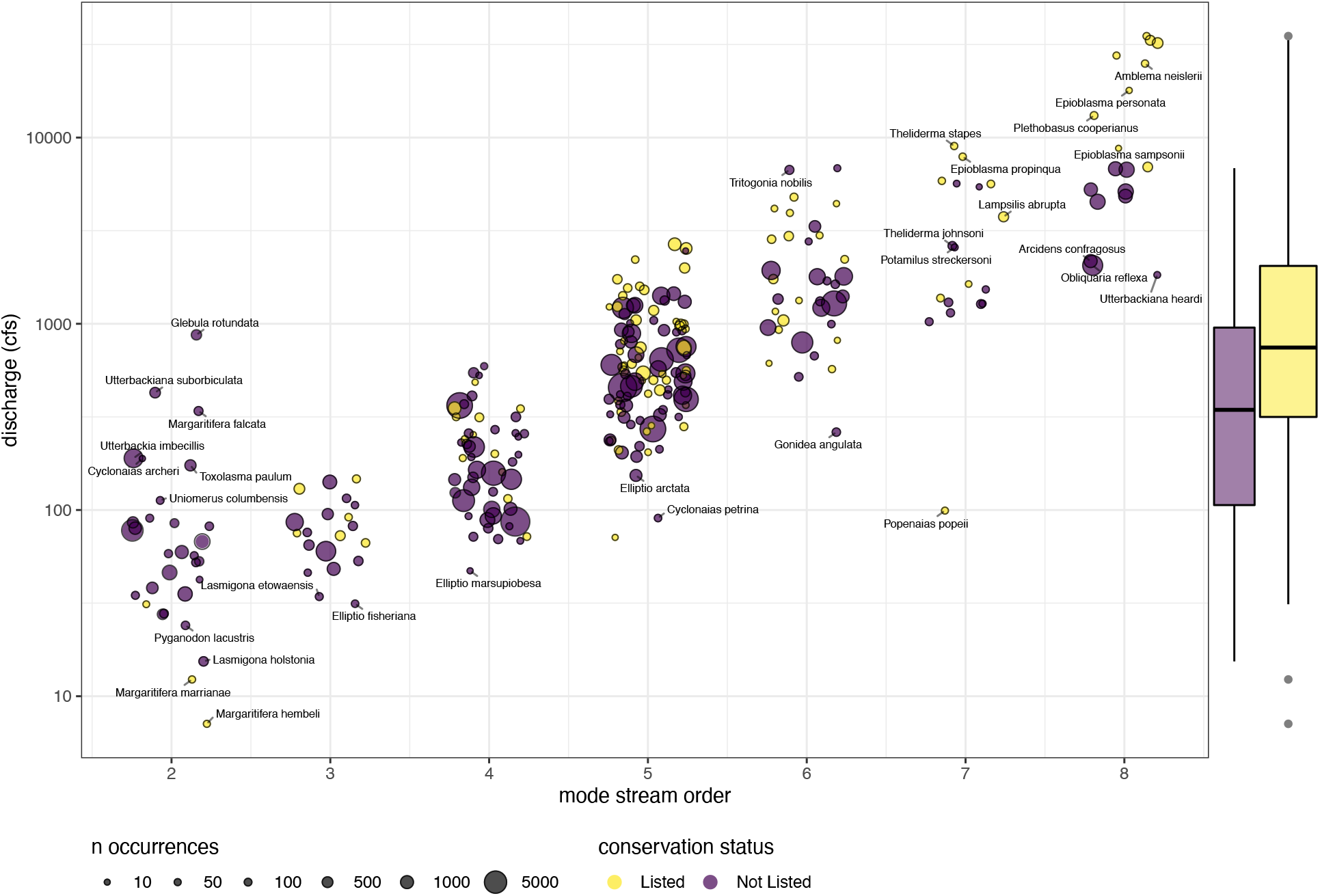
Jitter plot of each species mode stream order and median discharge preferences (log_10_ scale) with marginal box plot describing discharge variation within listed and non-listed species.

We assigned 241,484 non-flagged, species-level occurrences to 1,494 HUC8s. The number of occurrences per HUC8 varied dramatically from 1 occurrence (147 different HUC8s) to 7,940 occurrences (Upper Scioto) and had a median value of 29 occurrences per occupied HUC8 (Figure 6A). HUC8 species richness varied from 1 species (211 HUCs) to 92 species (Pickwick Lake, Tennessee River watershed) and had a median of 9 species (Figure 6B). Generalized patterns of US species richness recovered here are similar to previous treatments (Haag 2010, 2012), but estimates of species richness at the watershed level can vary substantially (Table 1). Estimates of percent change in HUC8 species diversity since median collection year varied from -95.6% (Blue Earth) to +71.2% (Canoochee) with a median percent change of -12.5 (Figure 7A). Occurrence poor HUC8s tended to have the greatest changes in species richness (Figure 7B). A summary of all HUC attributes is available in Appendix S2.

**Figure 6:**
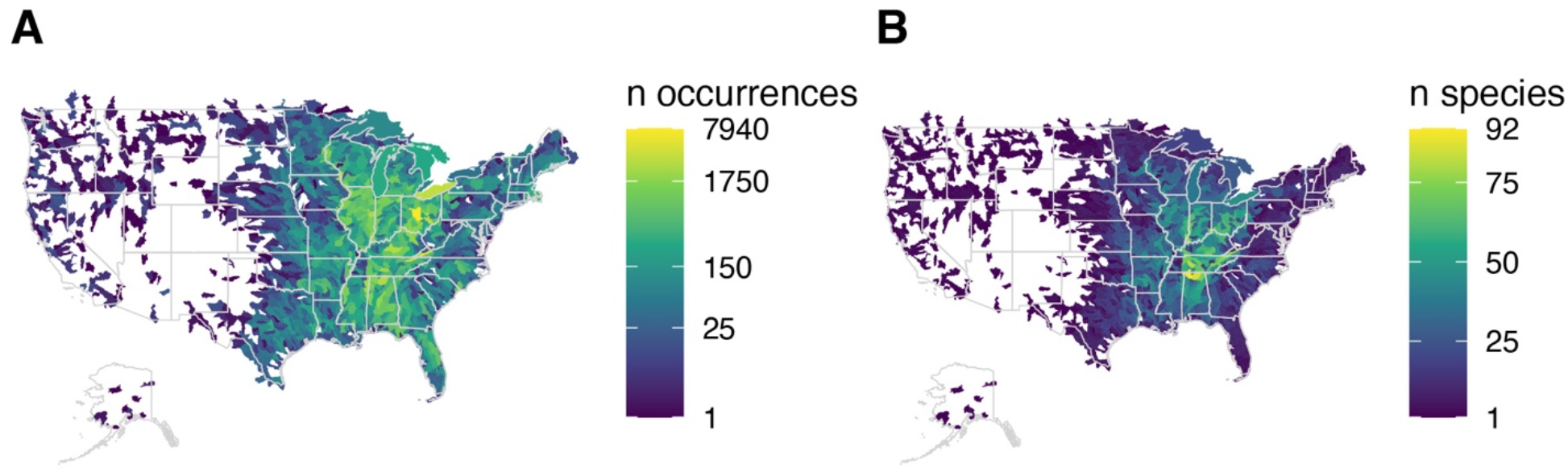
Choropleth describing the variation of georeferenced, species-level occurrences per HUC8 (A) and species richness per HUC8 (B).

**Figure 7:**
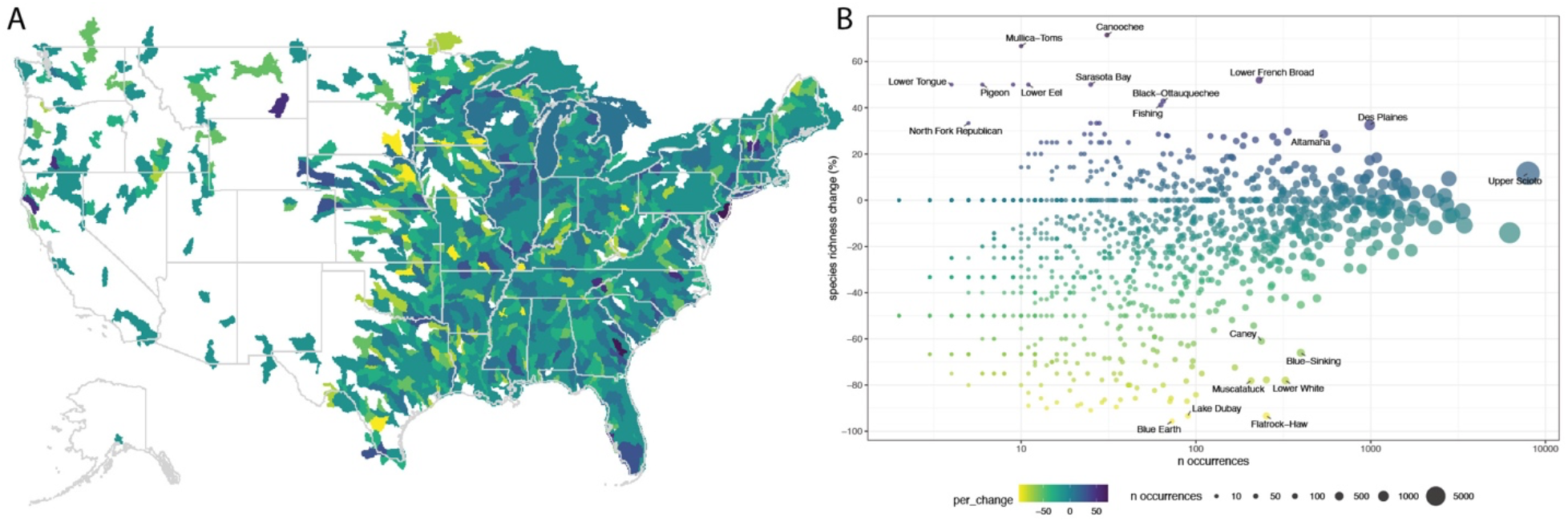
Choropleth describing the percent change in HUC8 species richness across the US (A), and bivariate of number of occurrences per HUC8 and percent change in HUC8 species richness(B).

**Table 1:**
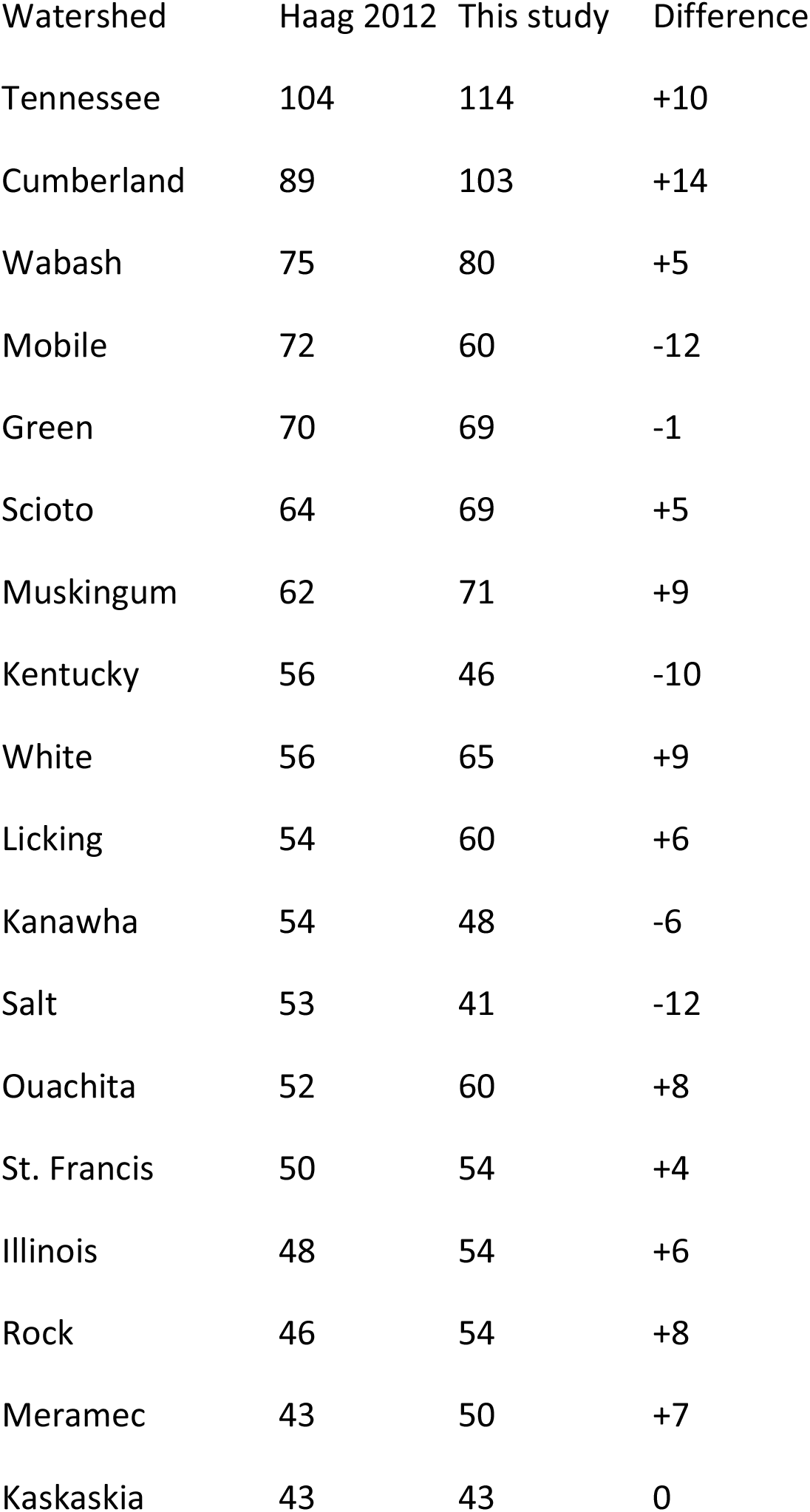
Comparison of species richness of 18 diverse watersheds in the US (Modified from Haag 2012)

## Discussion

### Inventory of Digitized US Freshwater Mussel Collections Data

With assistance from many collection managers and curators, we assembled an inventory of digitized US freshwater mussel collections data that includes 45 institutions and 410,665 records. This resource is a near comprehensive synthesis of digitized US freshwater occurrence data, but it is by no means complete. Additional specimen-based occurrences exist elsewhere, especially in natural history collections outside the US, as well as in the collections of federal and state wildlife agencies, academic research labs, and the undigitized portions of the included collections. Nevertheless, this synthesis is an important and timely step towards making US freshwater mussel collections data more centralized, standardized, and user-friendly for its diverse stakeholders.

Furthermore, we extended the utility of these records to better suit the needs of the freshwater mussel research and conservation community by standardizing taxonomic information (e.g. harmonizing >4,500 unique taxonomic identifications to 302 standardized species-level identifications), flagging potentially problematic records (e.g. duplicate records, misidentifications, inaccurate georeferences), integrating freshwater-specific spatial frameworks (e.g., hydrological units and stream segments), and joining records with various stream characteristics (e.g. stream order, annual mean velocity, annual mean discharge, and slope). This resource was used to explore temporal, taxonomic, ecological, and spatial patterns and make data-driven recommendations about ways to apply and improve freshwater mussel collections data as it relates to freshwater mussel research and conservation.

### Institutional Patterns

While there is significant specimen-based US freshwater mussel occurrence data from US institutions available via GBIF (∼189k records) and InvertEBase (∼210k records), this inventory provides a more comprehensive synthesis of US freshwater mussel collections data (∼410k records). This difference was driven by incorporating digitized data from numerous small collections (i.e., collections with <10k records) and one extremely large collection (i.e., OSUM; >70k records) that have not yet been integrated into GBIF or InvertEBase. Most freshwater mussel collections (∼70%) have less than 10,000 occurrences and most are very small in comparison to the largest collections. For example, there are more occurrences in the 6 largest freshwater mussel collections (221k) than in the remaining 39 collections assessed here (189k). The abundance of small freshwater mussel collections is characteristic of natural history collections at large (Monfils et al. 2020) and several studies have demonstrated the importance of small collections to understanding the diversity and distribution of various taxonomic groups (Sierwald et al. 2018; Cobb et al. 2019; Marsico et al. 2020).

The included institutions vary considerably in terms of their taxonomic and geographic coverage, as well as the degree of data completeness. Interestingly, no single institution had comprehensive species-level coverage of US freshwater mussel diversity and only in the aggregate is the entire mussel diversity of the country accounted for. Data completeness also varied substantially across the collections with several of the older freshwater mussel collections in the US clustering together around 50% of occurrences with year data and 80% of occurrences with georeferences. The numerous data-poor but historically important late 19th and early 20th century records at these institutions may prevent the inclusion of precise year data and georeferences. Every freshwater mussel collection has its strengths and weaknesses, and the identification of these areas is a useful first step towards more completely leveraging these data strengths and prioritizing collection improvements.

### Temporal Patterns

The US freshwater mussel research and conservation community has grown dramatically in the past couple of decades (Haag & Williams 2014; FMCS 2016). Presumably, this growth has resulted in an increase in some types of field work (e.g., sampling, monitoring, and surveying; see Smith et al. (2021)). Despite the expected (or locally demonstrated) growth in some field-based activities, freshwater mussel collecting (i.e., vouchering animals and depositing them in natural history museums) has decreased substantially in recent decades. Between 1970-2010 the average number of occurrences collected per year was 4,383, whereas between 2010-2020 the average number of mussel occurrences collected per year was 1,398 – a 68% decrease. The decrease in US freshwater mussel collecting is alarming and mirrors patterns observed in other faunas (Malaney & Cook 2018; Fischer et al. 2021; Rohwer et al. 2022). The decline of collecting is likely to be driven by numerous biological (e.g., declining mussel diversity, distribution, and abundance) and practical factors (e.g., collecting and collections are underfunded). Much of the decline observed here is driven by decreased collection of non-listed species and may be consistent with practical reasons having greater influence than biological ones.

Given the current resource limitations (e.g., staff, funding, space, supplies, permitting) it seems unlikely that individuals and institutions will return to the volume of collecting done in the latter half of the 20^th^ century. However, such heroic (and haphazard) collecting may not be necessary. By using existing collections data, we can develop more efficient, sustainable, and data-driven collecting strategies designed to increase impact over volume. For example, prioritizing collection of species and regions that are poorly represented in collection (i.e., illuminate dark taxa and watersheds) or avoid collection of species that have been recently collected from a particular area (i.e., reduce collection redundancy).

### Taxonomic patterns

US freshwater mussel taxonomy has changed considerably over the past 200 years and in the process has created numerous available names and subsequent combinations (Graf & Cummings 2021). As mussel systematics has evolved, many preferred specimen identifications have fallen in and out of prevailing usage, resulting in an overabundance of unique taxonomic identifications relative to the number of recognized species (>4.9k and 302, respectively). Standardizing taxonomic information within and across institutions was a critical first step towards assessing taxonomic patterns and most identifications were assigned to a valid taxon with high confidence using the GBIF Taxonomic Backbone.

While morphology-based identification of freshwater mussels can be difficult (e.g., atypical or very young specimens, vague or absent locality information, sympatric and cryptic species), most occurrences in our dataset (98%) were identified to the species-level. Less precise taxonomic identifications were frequently identified to the genus rank and were common in taxa with poorly characterized species boundaries (e.g., *Elliptio*). Determining taxonomic precision (i.e., the rank at which specimens are identified in the taxonomic hierarchy) is programmatically straightforward but determining identification accuracy is less so. However, we were able to programmatically flag and remove >10k occurrences that are likely to be misidentified or have erroneous locality data. This approach has helped to remove many erroneous occurrences, but physical examination of specimens will undoubtedly catch further errors.

All US freshwater mussel species are represented in this dataset by at least one occurrence except *Disconaias fimbriata*. That genus and species is recognized as part of the Recent US fauna (Williams et al. 2017) but based on the collections surveyed there is no voucher-based evidence that either occur in US fresh waters. Freshwater mussel occurrence data has a strong taxonomic bias with a continuum of occurrence-poor and occurrence-rich species. The identification of these taxonomic strengths and weaknesses should be useful in making data-driven research and collecting decisions like selecting a model study species or determining collecting priorities (e.g., occurrence-rich and occurrence-poor species, respectively).

### Ecological Attributes

Freshwater mussels of the US are an ecologically divergent assemblage with well documented variation in many species traits (e.g. fecundity, brooding morphology, parasitic larval morphology) (Haag 2012). In comparison many freshwater mussel ecological attributes are poorly quantified (e.g., aspects of range size and habitat preferences). Some of these ecological attributes are frequently used to characterize aspects of freshwater mussel diversity and ecology and include generalizations like “geographically widespread species” (Pfeiffer et al. 2018), “narrowly distributed species” (Strayer 2008), “headwater species” (Haag & Warren 1998), “large river specialists” (Haag 2012), “riffle-dwelling species” (Gagnon et al. 2004), and “slackwater associates” (Gagnon et al. 2006). Despite the ubiquity and usefulness of these generalizations, the underlying ecological attributes have not been meaningfully quantified across the assemblage. By bridging non-flagged, georeferenced, species-level records with freshwater-specific spatial frameworks, we were able to quantify aspects of several ecological attributes of 282 freshwater mussel species in the US. We discuss how two of these ecological attributes, range size and habitat use preferences, can be used to better understand patterns of freshwater mussel ecology and conservation.

Freshwater mussel geographic ranges are often clearly defined in the literature (although exceptions certainly exist) and usually align with discrete hydrological units (e.g., Mississippi River Drainage, Tennessee River subdrainage, Duck River watershed). Despite the clarity of freshwater mussel geographic ranges, the size of a species occupied habitat (i.e., AOO) is infrequently measured. This knowledge gap is unfortunate because AOO is a practically important ecological attribute in conservation biology as it is a useful predictor of extinction risk and sensitivity to climate change (Purvis et al. 2000; Mims et al. 2018; Chichorro et al. 2019).

The negative relationship between AOO and extinction risk (high AOO = low extinction risk, low AOO = high extinction risk) is strong to the point of being an operational criterion for determining conservation status (IUCN 2012). However range size is an infrequently used metric in conservation assessments associated with the Endangered Species Act; and unfortunately, range size was not included in any of the 15 freshwater mussel listing decisions between February 2011 to October 2014 (Smith-Hicks & Morrison 2021).

We bridged this important knowledge gap by estimating AOO for 282 of the 303 US freshwater mussel species. Given the relationship between AOO and extinction risk these estimates could be helpful to species status assessments and identifying conservation priorities. Listed species had a significantly smaller median AOO than non-listed species, consistent with the well-documented phenomenon that species with small range sizes tended to have greater extinction risk (Purvis et al. 2000; Mims et al. 2018; Chichorro et al. 2019). At the HUC8 level, natural history collections data is likely to underestimate AOO because of incomplete geographic sampling (i.e., false absences at HUC8 level). These underestimates may be particularly acute in occurrence-poor species, many of which tend to be listed species. At the same time, measuring AOO at the HUC8 level tends to overestimate range size of aquatic organisms because aquatic species are patchily distributed across that much larger hydrological unit (DeWeber & Wagner 2015; Frimpong et al. 2016; Mims et al. 2018). Range size estimates using greater spatial resolution (HUC10 or COMID) are possible with the data provided here and could be useful to creating more nuanced range maps and AOO estimates. We look forward to improvements of these range size estimates and insights as to how they vary as a function of other aspects of freshwater mussel natural history (e.g., body size, number of hosts, phylogeny).

An additional ecological attribute used as a primary criterion to evaluate conservation status is the change in a species AOO over time (IUCN - IUCN 2012; ESA - Smith et al. 2018). Natural history collections data is uniquely positioned to measure how species ranges have changed across time, but such insights are not without major challenges including significant temporal, taxonomic and spatial biases (Meyer et al. 2016; Kharouba et al. 2019; Davis et al. 2022). An additional complication of determining temporal changes is when subjective timeframes are used to define a species “historic” and “recent” distribution (e.g., pre- and post-1960 (Metcalfe-Smith et al. 1998); pre- and post-1990 (Blevins et al. 2017); pre- and post-1995 (Smith et al. 2021); pre- and post-2000 (Johnson et al. 2016)). Because collecting is taxonomically and temporally uneven subjectively defined timeframes can result in comparison of time frames with dramatically different collecting effort. Comparisons with greater collecting effort in the first timeframe will bias results towards decreasing AOO; greater collecting effort in the second time frame will bias results towards increasing AOO.

We attempted to standardize taxonomic and temporal biases by objectively splitting each species’ collection history into two timeframes with roughly equal collection effort (i.e., time frame defined by that median year of occurrences for that species). After accounting for these taxonomic and temporal biases, listed species experienced marginally greater decreases in AOO in comparison to non-listed species, but only when occurrence-poor species were excluded. There are many exceptions to this pattern which illustrate how collections can (and can’t) inform our understanding of species declines. For example, although *Epioblasma alhstedi* had the greatest decrease in AOO but is a non-listed species. Recommendations to list *E. alhstedi* date back >15 years (Jones et al 2006), to when the species was split out of the already federally endangered species *E. capsaeformis*. On the other hand, *Paravaspina stensteiana* is listed as federally endangered but has an extremely high estimated increase in AOO. This exception may be due to the very few records used to estimate percent change in AOO (n=21) or an increase in scientific effort following its listing (Smith-Hicks & Morrison 2021).

Percent change in AOO (and other attributes measured here) are volatile when estimated for occurrence-poor species and this fact should be carefully considered when drawing inferences. Our method of accounting for temporal variation in collecting effort minimizes changes in AOO and is designed to be a conservative estimate of changes in AOO (at least in occurrence-rich taxa). Integrating specimen-based collections data with additional data sources, especially the abundance of state and federal survey data, via integrated modeling could be useful to improving these estimates (Davis et al. 2022).

Habitat preferences are a frequently mentioned ecological attribute in the literature and is obvious in riverscapes at large and small scales (entire drainages to riffle-pool sequences)(Strayer 2008; Haag 2012). However, many aspects of habitat preference have not yet been rigorously quantified (e.g., stream order, discharge, velocity, slope). For example, various freshwater mussel species have been referred to as “headwater species’’ or “large-river species” (Haag & Warren 1998; Gagnon et al. 2006; Pandolfi et al. 2022) but those characterizations remain amorphous and anecdotal. Our quantification of stream order and discharge preferences reflect these previously hypothesized generalizations and appear to be ecologically relevant. Taxa often considered to be headwater species’ (e.g., several members of the Margaritiferinae, *Lasmigona, Elliptio*) were estimated to have lower stream order preferences and lower discharge preferences (e.g. 2-3 and <100 cfs, respectively) in comparison to taxa considered to be large-river species (e.g. several members of the Amblemini, *Potamilus, Obovaria*) which tended to have higher stream order preferences and higher discharge preferences (e.g. 4-8 and >300 cfs, respectively). Despite numerous unaccounted for misidentifications and inaccurate georeferences/stream segment assignments, which are certain to add noise, our species-specific estimates support previous hypotheses and provide a numerical framework to quantify habitat preference and explore its role in mussel evolution, ecology and conservation. We also observed that listed species had larger stream order and discharge preferences than non-listed species. The recovered relationship between habitat preference and conservation status is consistent with the systematic destruction of US riverine ecosystems via damming and channelization in the middle of the 20^th^ century, which primarily affected large rivers and their obligate freshwater mussel assemblages (i.e. “first extinction wave” – Haag 2009, 2012).

### Spatial Patterns

The entire US freshwater mussel research and conservation community relies heavily on a remarkable compendium of regional mussel biodiversity atlases (e.g., Williams et al. 2008; Watters et al. 2009; Haag & Cicerello 2016). These invaluable resources provide intensely researched and expertly curated biogeographic information focused on species of geopolitical areas (primarily US states), but no nation-wide atlas is yet available. This inventory is a useful start to that end. While the level and type of curation in this dataset is distinctly different from previous published regional atlases (programmatic flagging vs physically examining specimens), this resource benefits from being nationally comprehensive and based on explicit and easily accessible records.

By integrating natural history collection data into spatial frameworks designed for freshwater resource management, we provide the first summary of how occurrence data varies across hydrological units of the US and enables conservation biologists and wildlife managers with more accessible and practically useful biogeographic information. Leveraging this resource, we were able to quantify how collecting effort and species richness varies across five standardized hydrological levels (HUC2-HUC10). While this approach is useful, our analysis of spatial distribution only includes the georeferenced occurrences and ignores ∼30% of all US freshwater mussel occurrences. Identification of these geographic and data strengths and weaknesses will be useful for directing field work towards under-collected regions and improving spatial data completeness.

Our occurrence-based estimates of species richness across hydrological units were greatest in the Tennessee and Ohio Rivers, consistent with previous large scale biogeographic assessments of US freshwater mussels (Haag 2010, 2012). However, at smaller scales our estimates can be quite different from previous estimates. Differences in species richness between sources should be investigated further and may be products of differing taxonomy, misidentifications, incorrect geographic information, or real biological changes in species diversity.

While accounting for temporal collecting bias, we observed widespread increases and decreases in HUC species richness. Generally, hydrological units in the western US and the western edge of the Mississippian River region had the largest losses in species richness. This pattern could have resulted from under sampling in the region or true losses of species diversity via marginal range contraction, or a combination of the two. On average, species diversity at the HUC8 level has decreased 17.8%, although occurrence poor HUCs had volatile estimates. It remains unclear if these changes in HUC species richness are an indication of changes in collection effort or true losses of local biodiversity, but in either case, this analysis identifies areas to potentially investigate as mussel decline continues.

This synthesis has brought to light a more complete picture of digitized US freshwater mussel collections data and illuminated various institutional, temporal, taxonomic, spatial and ecological patterns. While we have attempted to flag problematic occurrences, many errors remain unaccounted for. Despite that certainty, clear patterns have emerged that are consistent with previous biogeographic scenarios, ecological hypotheses, and conservation assessments. We are hopeful that this resource will serve as a useful starting point for various aspects of taxon- and habitat-based conservation efforts (e.g., taxonomic revisions, species status assessments, listing decisions, recovery plans), as well as be used by natural history collections to make data driven decisions about how to improve the data they curate. Updating existing records and integrating new collections into more centralized, resourced, and frequently updated biodiversity aggregators like GBIF and InvertEBase should be a pressing goal. Beyond increasing the proportion of georeferenced occurrences there is a clear need for more standardized, accurate, and nuanced geodetic data (e.g., including spatial uncertainty).

Finally, the attrition of freshwater mussel collecting in the US in recent decades has the potential to have lasting effects in research and conservation, but many institutions may not have the resources available to support more collecting. Our collecting strategies must evolve. If collecting less is a requirement, we should collect more strategically. We have outlined several ways in which this resource could be used to determine when, what, and where to collect vouchers (e.g., occurrence-poor species, occurrence-poor HUCs, new or outdated HUC-occurrence). This type of data-driven collecting has the potential to decrease investment but increase return. The freshwater mussel conservation community has a strong record of coordinating successful national strategies but collecting has not yet been addressed in any of these visions (NNMC 1998; Haag & Williams 2014; FMCS 2016). Integrating data-driven collection practices into these national strategies have the potential to improve efforts aimed at conserving one of the world’s most imperiled regional faunas.

## Acknowledgements

We are grateful to the collection mangers and curators that helped us to create this resource: K. Kocot (ALMNH), L. Berniker (AMNH), G. Rosenberg (ANSP), M. Ganglof (APPZ), J. Harris (ASUMZ), N. Noor (AUMNH), J. Leah (BMSM), M. Turk (BSNS), D. Roberts (CHAS), T. Pearce (CM), C. Brennan (CMNH), A. Kittle (DMNH), C. Carter (DMNS), D. Hayes (EKU), J. Gerber (FMNH), J. Wares (GMNH), R. Vinsel (INHS), J. Sheff (INSM), A. Simons (JFBM), K. Jensen (KU), L. Groves (LACM), A. Lewis (LSUS), A. Baldinger (MCZ), G. Dinkins(MMNHC), R. Ellwanger (MMNS), H. Farrington (MNHSC), J. Zaspel (MPM), A. Bogan (NCSM), D. Mayer (NYSM), A. Franzen (OBS), J. Braun (OMNH), N. Shoobs (OSUM), K. Morton (PMNS), A. Benedict (SMBU), L. Appleton (TNHC), M. Suter (UAFMC), B. Chalifour (UCM), J. Slapsinsky (UF), T. Lee (UMMZ), K. Ahfield (USNM), V. Zhuang (UTEP), M. Frey (UWBM), L. Monahan (UWZM), J. Harris (VMNH), and L. Rojas (YPM). The code available at D. MacGuigan’s “Hidden Rivers of the United States” project inspired our programmatic out-of-range flagging approach. We are grateful to C. Atkinson and G. Hopper for their suggestions to include estimates of each species minimum, maximum, and centroid georeferences.

## Supporting information

Additional supporting information may be found in the online version of the article at the publisher’s website. All data and code necessary for reproduction are available at https://doi.org/10.5061/dryad.c2fqz61cg.

Appendix S1: Code to standardize, process, analyze, and visualize collections data

Appendix S2: All .csv data files

Appendix S3: Species range shapefiles for programmatic flagging

## References

Blevins E, Jepsen S, Box JB, Nez D, Howard J, Maine A, O’Brien C. 2017. Extinction risk of western North American freshwater mussels: Anodonta nuttalliana, the Anodonta oregonensis/kennerlyi clade, Gonidea angulata, and Margaritifera falcata. Freshwater Mollusk Biology and Conservation 20:71–88.

Campbell Grant EH. 2015. Please don’t misuse the museum:’declines’ may be statistical. Global Change Biology 21:1018–1024.

Chamberlain S, Ram K, Barve V, Mcglinn D, Chamberlain MS. 2017. Package ‘rgbif’. Interface to the Global Biodiversity Information Facility ‘API 5:0.9.

Chang W, Cheng J, Allaire J, Sievert C, Schloerke B, Xie Y, Allen J, McPherson J, Dipert A, Borges B. 2022. shiny: Web Application Framework for R. R package version 1.7.2. <https://CRAN.R-project.org/package=shiny>.

Chichorro F, Juslén A, Cardoso P. 2019. A review of the relation between species traits and extinction risk. Biological Conservation 237:220–229.

Cobb NS, Gall LF, Zaspel JM, Dowdy NJ, McCabe LM, Kawahara AY. 2019. Assessment of North American arthropod collections: Prospects and challenges for addressing biodiversity research. PeerJ 7:e8086.

NNMC. 1998. National strategy for the conservation of native freshwater mussels. Journal of Shellfish Resources 17:1419–1428.

Davis CL, Guralnick RP, Zipkin EF. 2022. Challenges and opportunities for using natural history collections to estimate insect population trends. Journal of Animal Ecology.

DeWeber JT, Wagner T. 2015. Predicting brook trout occurrence in stream reaches throughout their native range in the eastern United States. Transactions of the American Fisheries Society 144:11–24.

Fischer EE, Cobb NS, Kawahara AY, Zaspel JM, Cognato AI. 2021. Decline of amateur lepidoptera collectors threatens the future of specimen-based research. BioScience 71:396–404.

FMCS. 2021. The 2021 checklist of freshwater bivalves (Mollusca: Bivalvia: Unionida) of the United States and Canada. Considered and approved by the Bivalve Names Subcommittee December 2020.

Frimpong EA, Huang J, Liang Y. 2016. IchthyMaps: A database of historical distributions of freshwater fishes of the United States. Fisheries 41:590–599.

Gagnon P, Michener W, Freeman M, Box JB. 2006. Unionid habitat and assemblage composition in coastal plain tributaries of Flint River (Georgia). Southeastern Naturalist 5:31–52.

Gagnon PM, Golladay SW, Michener WK, Freeman MC. 2004. Drought responses of freshwater mussels (Unionidae) in coastal plain tributaries of the Flint River basin, Georgia. Journal of Freshwater Ecology 19:667–679.

GBIF Secretariat. 2021. GBIF Backbone Taxonomy. Checklist dataset https://doi.org/10.15468/39omei accessed via GBIF.org on 2022-09-16.

Graf DL, Cummings KS. 2021. A ‘big data’approach to global freshwater mussel diversity (Bivalvia: Unionoida), with an updated checklist of genera and species. Journal of Molluscan Studies 87:eyaa034.

Haag WR. 2009. Past and future patterns of freshwater mussel extinctions in North America during the Holocene. Holocene extinctions: 107–128.

Haag WR. 2010. A hierarchical classification of freshwater mussel diversity in North America. Journal of Biogeography 37:12–26.

Haag WR 2012. North American freshwater mussels: natural history, ecology, and conservation. Cambridge University Press.

Haag WR. 2019. Reassessing enigmatic mussel declines in the United States. Freshwater Mollusk Biology and Conservation 22:43–60.

Haag WR, Cicerello RR 2016. A distributional atlas of the freshwater mussels of Kentucky. Kentucky State Nature Preserves Commission.

Haag WR, Warren J, Melvin L. 1998. Role of ecological factors and reproductive strategies in structuring freshwater mussel communities. Canadian Journal of Fisheries and Aquatic Sciences 55:297–306.

Haag WR, Williams JD. 2014. Biodiversity on the brink: an assessment of conservation strategies for North American freshwater mussels. Hydrobiologia 735:45–60.

Hedrick BP, Heberling JM, Meineke EK, Turner KG, Grassa CJ, Park DS, Kennedy J, Clarke JA, Cook JA, Blackburn DC. 2020. Digitization and the future of natural history collections. BioScience 70:243–251.

IUCN. 2012. IUCN Red List Categories and Criteria: Version 3.1. Second edition. Gland, Switzerland and Cambridge, UK: IUCN. iv + 32pp.

Johnson NA, McLeod JM, Holcomb J, Rowe M, Williams JD. 2016. Early life history and spatiotemporal changes in distribution of the rediscovered Suwannee moccasinshell Medionidus walkeri (Bivalvia: Unionidae). Endangered Species Research 31:163–175.

Kharouba HM, Lewthwaite JM, Guralnick R, Kerr JT, Vellend M. 2019. Using insect natural history collections to study global change impacts: challenges and opportunities. Philosophical Transactions of the Royal Society B 374:20170405.

Malaney JL, Cook JA. 2018. A perfect storm for mammalogy: declining sample availability in a period of rapid environmental degradation. Journal of Mammalogy 99:773–788.

Marsico TD, Krimmel ER, Carter JR, Gillespie EL, Lowe PD, McCauley R, Morris AB, Nelson G, Smith M, Soteropoulos DL. 2020. Small herbaria contribute unique biogeographic records to county, locality, and temporal scales. American journal of botany 107:1577–1587.

McKay L, Bondelid T, Dewald T, Johnston J, Moore R, Rea A. 2012. NHDPlus Version 2: User Guide.

Metcalfe-Smith JL, Staton SK, Mackie GL, Lane NM. 1998. Changes in the biodiversity of freshwater mussels in the Canadian waters of the lower Great Lakes drainage basin over the past 140 years. Journal of Great Lakes Research 24:845–858.

Meyer C, Weigelt P, Kreft H. 2016. Multidimensional biases, gaps and uncertainties in global plant occurrence information. Ecology letters 19:992–1006.

Mims MC, Olson DH, Pilliod DS, Dunham JB. 2018. Functional and geographic components of risk for climate sensitive vertebrates in the Pacific Northwest, USA. Biological Conservation 228:183–194.

Monfils AK, Krimmel ER, Bates JM, Bauer JE, Belitz MW, Cahill BC, Caywood AM, Cobb NS, Colby JB, Ellis SA. 2020. Regional collections are an essential component of biodiversity research infrastructure. BioScience 70:1045–1047.

Moore RB, McKay LD, Rea AH, Bondelid TR, Price CV, Dewald TG, Johnston CM. 2019. User’s guide for the national hydrography dataset plus (NHDPlus) high resolution. Report. Survey USGS, Reston, VA.

Nelson G, Ellis S. 2019. The history and impact of digitization and digital data mobilization on biodiversity research. Philosophical Transactions of the Royal Society B 374:20170391.

Pandolfi GS, Mays JW, Gangloff MM. 2022. Riparian land-use and in-stream habitat predict the distribution of a critically endangered freshwater mussel. Hydrobiologia 849:1763–1776.

Pebesma EJ. 2018. Simple features for R: standardized support for spatial vector data. R J. 10:439.

Pfeiffer JM, Sharpe AE, Johnson NA, Emery KF, Page LM. 2018. Molecular phylogeny of the Nearctic and Mesoamerican freshwater mussel genus Megalonaias. Hydrobiologia 811:139–151.

Pursifull S, Holcomb J, Rowe M, Williams JD, Wisniewski JM. 2021. Status of freshwater mussels in the Ochlockonee River Basin of Georgia and Florida. Southeastern Naturalist 20:1–19.

Purvis A, Gittleman JL, Cowlishaw G, Mace GM. 2000. Predicting extinction risk in declining species. Proceedings of the royal society of London. Series B: Biological Sciences 267:1947–1952.

Rohwer VG, Rohwer Y, Dillman CB. 2022. Declining growth of natural history collections fails future generations. PLoS Biology 20:e3001613.

Service USFaW. 50 CFR 17.11.

Sierwald P, Bieler R, Shea EK, Rosenberg G. 2018. Mobilizing mollusks: Status update on mollusk collections in the USA and Canada. American Malacological Bulletin 36:177–214.

Smith CH, Johnson NA, Inoue K, Doyle RD, Randklev CR. 2019. Integrative taxonomy reveals a new species of freshwater mussel, Potamilus streckersoni sp. nov.(Bivalvia: Unionidae): implications for conservation and management. Systematics and Biodiversity 17:331–348.

Smith CH, Johnson NA, Robertson CR, Doyle RD, Randklev CR. 2021. Establishing conservation units to promote recovery of two threatened freshwater mussel species (Bivalvia: Unionida: Potamilus). Ecology and Evolution 11:11102–11122.

Smith DR, Allan NL, McGowan CP, Szymanski JA, Oetker SR, Bell HM. 2018. Development of a species status assessment process for decisions under the US Endangered Species Act. Journal of Fish and Wildlife Management 9:302–320.

Smith-Hicks KN, Morrison ML. 2021. Factors Associated with Listing Decisions under the US Endangered Species Act. Environmental Management 67:563–573.

FMCS. 2016. A national strategy for the conservation of native freshwater mollusks. Freshwater Mollusk Biology and Conservation 19:1–21.

Strayer DL 2008. Freshwater mussel ecology: a multifactor approach to distribution and abundance. Univ of California Press.

Team RC. 2022. R: A language and environment for statistical computing. R Foundation for Statistical Computing, Vienna, Austria. https://www.R-project.org/.

USGS. 2020. U.S. Geological Survey. National Watershed Boundary Dataset (ver. USGS National Watershed Boundary Dataset in FileGDB 10.1 format (published 20200701), accessed February 28, 2021 at https://www.usgs.gov/national-hydrography/access-national-hydrography-products.

USFWS Species Data Explorer, Available from https://ecos.fws.gov/ecp/report/adhocDocumentation?catalogId=species&reportId=species (accessed September 13 2022).

Watters GT, Hoggarth MA, Stansberry DH 2009. The freshwater mussels of Ohio. The Ohio State University Press.

Williams JD, Bogan AE, Butler RS, Cummings KS, Garner JT, Harris JL, Johnson NA, Watters GT. 2017. A revised list of the freshwater mussels (Mollusca: Bivalvia: Unionida) of the United States and Canada. Freshwater Mollusk Biology and Conservation 20:33–58.

Williams JD, Bogan AE, Garner JT 2008. Freshwater mussels of Alabama and the Mobile basin in Georgia, Mississippi, and Tennessee. University of Alabama Press.

